# Multi-Experiment Nonlinear Mixed Effect Modeling of Single-Cell Translation Kinetics after Transfection

**DOI:** 10.1101/285478

**Authors:** F. Fröhlich, A. Reiser, L. Fink, D. Woschée, T. Ligon, F. J. Theis, J. O. Rädler, J. Hasenauer

## Abstract

Single-cell time-lapse studies have advanced the quantitative understanding of cell-to-cell variability. However, as the information content of individual experiments is limited, methods to integrate data collected under different conditions are required.

Here we present a multi-experiment nonlinear mixed effect modeling approach for mechanistic pathway models, which allows the integration of multiple single-cell perturbation experiments. We apply this approach to the translation of green fluorescent protein after transfection using a massively parallel read-out of micropatterned single-cell arrays. We demonstrate that the integration of data from perturbation experiments allows the robust reconstruction of cell-to-cell variability, i.e., parameter densities, while each individual experiment provides insufficient information. Indeed, we show that the integration of the datasets on the population level also improves the estimates for individual cells by breaking symmetries, although each of them is only measured in one experiment. Moreover, we confirmed that the suggested approach is robust with respect to batch effects across experimental replicates and can provide mechanistic insights into the nature of batch effects. We anticipate that the proposed multi-experiment nonlinear mixed effect modeling approach will serve as a basis for the analysis of cellular heterogeneity in single-cell dynamics.

## Introduction

After decades of investigation, it is well known that even isogenic cell populations show molecular and phenotypic differences at the single-cell level (Altschuler & Wu 2010; Elowitz et al. 2002). Sources of cell-to-cell variability include noisy cellular processes (Elowitz et al. 2002), differences in cell cycle state (Buettner et al. 2015), the history of individual cells (Spencer et al. 2009), as well as spatio-temporal differences of the cell’s environment (Snijder & Pelkmans 2011). Methods such as mass cytometry (Bodenmiller et al. 2012) or single-cell RNA sequencing (Angerer et al. 2017) can provide highly multiplexed snapshots of cell-to-cell variability in thousands to millions of cells. Complementarily, time-lapse microscopy allows for the time-resolved measurement of cell-to-cell variability in the dynamic response of cells (Muzzey & van Oudenaarden 2009; Locke & Elowitz 2009). Recently, in order to improve the high-throughput capability of single-cell time-lapse studies, microstructured arrays (Muzzey & van Oudenaarden 2009) or microfluidic devices (Uhlendorf et al. 2012) are used to restrict cells in their movement, enabling automated acquisition of single-cell fluorescence trajectories over time.

Single-cell technologies already facilitated many novel insights, ranging from the analysis of population structures (Bodenmiller et al. 2012; Buettner et al. 2015) over the assessment of developmental trajectories (Navin et al. 2011; Haghverdi et al. 2016) to mechanistic insights into causal differences (Elowitz et al. 2002; Spencer & Sorger 2011; Hasenauer et al. 2014; Loos et al. 2017). To gain mechanistic insights, many studies use ordinary differential equation (ODE) models (Chen et al. 1998; Kühn et al. 2009; Klipp et al. 2005; Kitano 2002).

In this spirit, earlier studies have analyzed time-lapse microscopy measurements of single-cells after transfection with synthetic mRNA to assess mRNA lifetime (Leonhardt et al. 2014). mRNA lifetime is of fundamental interest to basic science, as it is a key parameter in many gene regulatory processes. Moreover, transient transfection of synthetic mRNA is relevant for biomedical applications, as it enables treatment of diseases via the targeted expression of proteins (Yamamoto et al. 2009; Kreiter et al. 2011). Hence, a good understanding and control of the expression dynamics of therapeutic proteins is essential for treatment design (Sahin et al. 2014). Yet, inference of quantitative estimates from single-cell experiments is model dependent and only insofar meaningful as our mechanistic understanding of many basic cellular processes, including transcription and translation, is sufficiently accurate. The model parameters can be estimated from single-cell time-lapse microscopy measurements using two different approaches:

**(I)** The standard two-stage approach (STS) estimates single-cell parameters and population distribution parameters sequentially (Almquist et al. 2015; Karlsson et al. 2015). First, parameters for every single cell are estimated independently by fitting an ODE to the respective trajectory. Then, a population-wide parameter distribution is reconstructed according to the single-cell parameter estimates. The STS approach enjoys great popularity (Leonhardt et al. 2014; Kalita et al. 2011; Karlsson et al. 2015; Almquist et al. 2015), because it is easy to implement, as many methods and tools developed for bulk data can be applied. However, the STS approach fails to distinguish between cell-to-cell variability and uncertainty of the estimated single-cell parameters, resulting in the overestimation of cell-to-cell variability (Sheiner & Beal 1983). This impairs applicability of the STS approach in settings with high experimental noise and sparse observations (Karlsson et al. 2015).

**(II)** In contrast, the non-linear mixed effect (NLME) approach (Pinheiro 1994) estimates single-cell parameters and population distribution parameters simultaneously. The single-cell parameters are considered as latent variables, which are constrained by the population distribution. The implementation of the NLME approach is more involved (Wang 2007; Kuhn & Lavielle 2005; Beal & Sheiner 1980) and it’s application computationally more intensive. Originally developed in pharmacology (Beal & Sheiner 1980), the NLME approach has recently risen in popularity for the analysis of single-cell data (Zechner et al. 2014; Karlsson et al. 2015; Almquist et al. 2015; Llamosi et al. 2016). It has been reported that NLME is more robust than STS in settings with large parameter uncertainty, as it reduces uncertainty (Sheiner & Beal 1983; Karlsson et al. 2015) and removes estimation bias (Almquist et al. 2015).

The NLME approach has several advantages over the STS approach when single-cell parameters have poor practical identifiable (Sheiner & Beal 1983; Karlsson et al. 2015), i.e., when the amount or noisiness of the data prohibits reliable parameter estimation. However, structural non-identifiability (Chis et al. 2011) of single-cell parameters is problematic for the STS as well as for the NMLE approach. Structural non-identifiabilities, meaning that the reliable parameter estimation is impossible due to model structure (vector field and observable), of single-cell parameters may lead to structural non-identifiability of population distribution parameters (Lavielle & Aarons 2016) and thus prohibit the reliable estimation of cell-to-cell variability. For bulk data, such structural non-identifiabilities can be resolved by considering perturbation experiments (Flassig & Sundmacher 2012). For single-cell data, it is unclear how the consideration of perturbation experiments affects non-identifiability for the STS and NLME approach.

Previous studies have shown that the single-cell degradation rates of mRNAs and proteins are structurally non-identifiable when considering time-lapse microscopy measurements for a single protein (Ferizi et al. 2015). This also holds for the respective population average parameters, as long as no further assumptions are made (Sahin et al. 2014). For this application, the structural non-identifiability is particularly problematic, as it impedes the reliable estimation of the mRNA lifetime, a key parameters of interest.

In this study, we address this problem by extending NLME to a multi-experiment setting, allowing the integration of single-cell perturbation experiments, which is not possible for the STS approach. We apply the method to study the fluorescence trajectories of hundreds of individual cells after transfection with mRNA encoding for eGFP. In contrast to previous studies, experiments were carried out twofold using two distinct variants of eGFP that differ in their protein lifetime. By analyzing single-cell trajectories from both experiments in a consistent nonlinear mixed effect modeling approach, we demonstrate that both protein and mRNA degradation rates can be uniquely identified. Furthermore, we assess the use of extended models for translation, including enzymatic degradation as well as ribosomally limited translation and find evidence for ribosomal rate-limitation of the translation process. Moreover, we show that the developed approach enables the robust estimation of population parameters, despite the presence of batch effects.

## Results

### Single-cell time-lapse experiments reveal a large heterogeneity of the fluorescent reporter protein expression after mRNA transfection

To obtain high-quality data of single-cell transfection-translation dynamics, we combined micropatterned protein arrays (Fig. 1a) with scanning time-lapse microscopy of multiple positions over a duration of 30 h (Fig. 1b) using a tailored perfusion tubing system. The perfusion tubing system allows for mRNA transfection during the time-lapse measurement. The cells were incubated with mRNA lipoplexes (Lipofectamine2000™) during the first hour of the measurement and washed with cell culture medium afterwards to limit the time window for lipoplex uptake. After a cell successfully internalizes mRNA lipoplexes, the corresponding mRNA molecules are translated into fluorescent proteins. This translation processes can be described by biochemical rate equations (Fig. 1c) (Leonhardt et al. 2014). The use of tubing systems allows for the observation of the translation kinetics of single cells right after adding mRNA lipoplexes (Fig. 1d), which was not possible in previous studies. The protein expression dynamics of single cells were measured by integration over the fluorescence intensities of successfully confined and transfected cells (at least 500 cells per mRNA construct and experiment). The analysis of the trajectories revealed substantial cell-to-cell variability in the amplitude and timing of expression. The collected data are provided in the Supplementary File S1.

**Figure 1:**
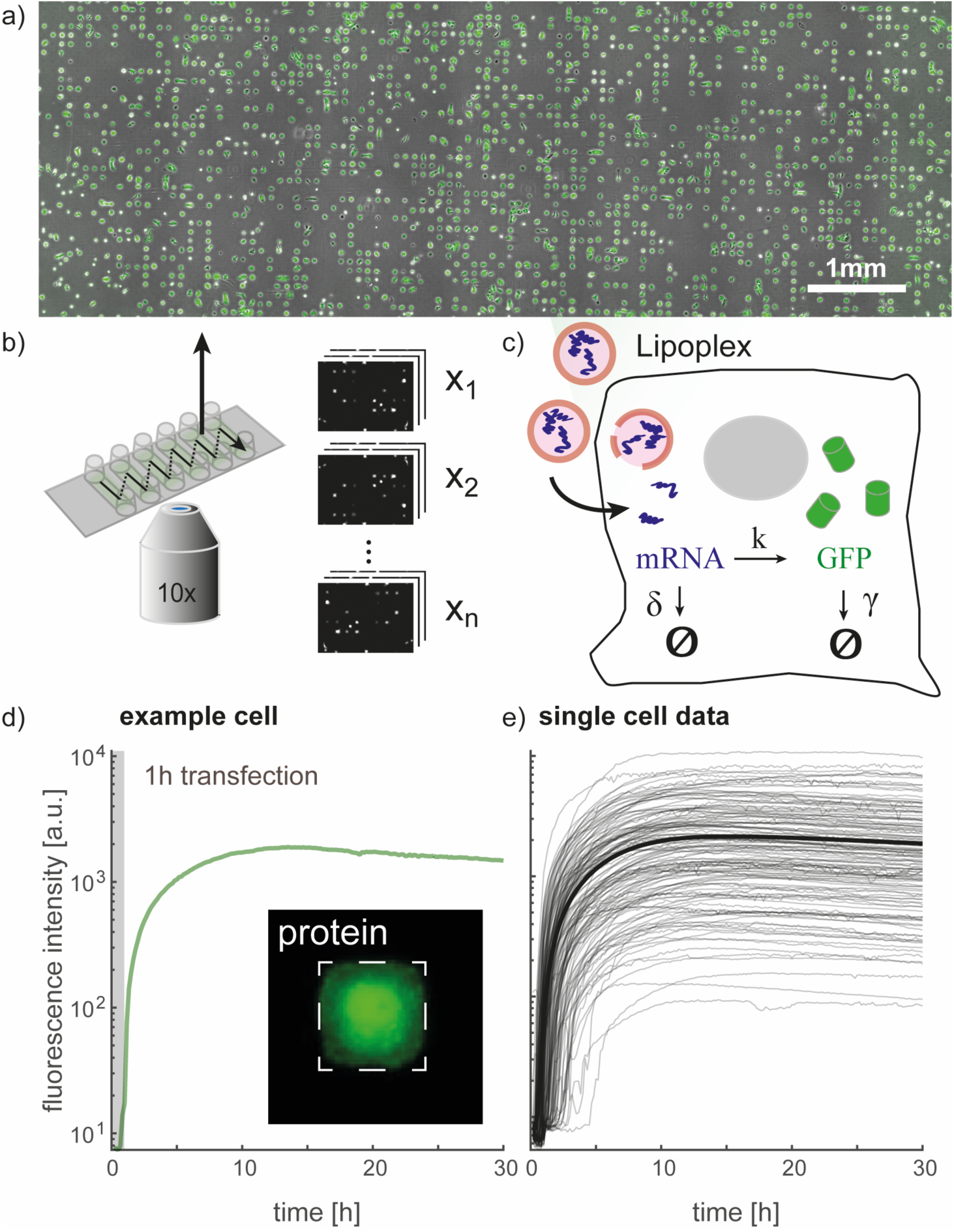
Single-cell translation assay for highly parallel readout of reporter protein expression kinetics after mRNA transfection. a) Micropatterned protein arrays are used for highly parallel readout of single-cell kinetics on standardized protein adhesion spots, which enables the observation of thousands of cells over a long time period. The microscopy image shows the micropatterned area of one channel with cells expressing eGFP. b) Schematics of the six-channel sample holder and the scanning time-lapse acquisition mode. Stacks of images from individual panels are depicted on the right. c) Schematic illustration of the transfection process using mRNA containing lipoplexes. The mRNA, which is released into the cytosol, is translated into a fluorescent reporter protein. The translation dynamics are modeled by biochemical rate equations. d) Single-cell eGFP expression is measured by integration over the fluorescence intensity. The zoom-in shows one eGFP-expressing cell confined on a fibronectin square (dashed square). The recording of protein expression begins by adding the mRNA lipoplexes, which are incubated for 1h. e) A subset of the single-cell trajectories of eGFP expressing cells shows the heterogeneity within the population. The thick black trajectory corresponds to the mean protein expression dynamic.

### Single-Experiment Single-Cell Measurements are Insufficient for Estimation of Protein Translation Parameters

For an in-depth analysis of the collected single-cell trajectories, we employed mathematical modeling. The process model we used is based on the established two-stage model of gene expression (Chen et al. 1998) and describes the concentration of mRNA and GFP molecules over time:

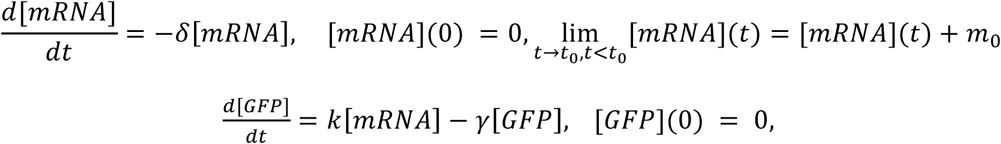

where *k* is the translation rate per mRNA, *m*_0_ is the amount of transfected mRNA entering the cell at time point *t*_0_, *δ* is the mRNA degradation rate and *γ* is the protein degradation rate. The measured fluorescence *y* is assumed as the sum of a signal proportional to the amount of eGFP (with scaling parameter *scale*) plus background fluorescence (*offset*). To facilitate the use of an additive error model, we consider the logarithm of the fluorescence intensity

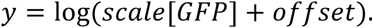

Not all of these parameters are structurally identifiable. Therefore, we reduced the model to a set of parameters *θ* that consists of products of the original parameters (see Supplementary Information, Section *Mathematical Models For GFP Translation*).

Following the literature (Almquist et al. 2015; Karlsson et al. 2015), we estimated all parameters using the STS approach (Fig. 2a). The parameters of individual cells were inferred from measured single-cell trajectories using a maximum likelihood method (stage 1) and then the distribution of parameters across the cell population was assessed (stage 2). The single-cell trajectories indicate pronounced cell-to-cell variability (Fig. 2b), which is reasonably well captured in the individual single-cell model fits (Fig. 2c). This pronounced cell-to-cell variability is in agreement with previous studies suggesting stochasticity of mRNA uptake (Ligon et al. 2014) and limited enzyme abundances as well as inhomogeneous spatial distribution within cells, e.g. for ribosomes (Liu et al. 2016).

**Figure 2.**
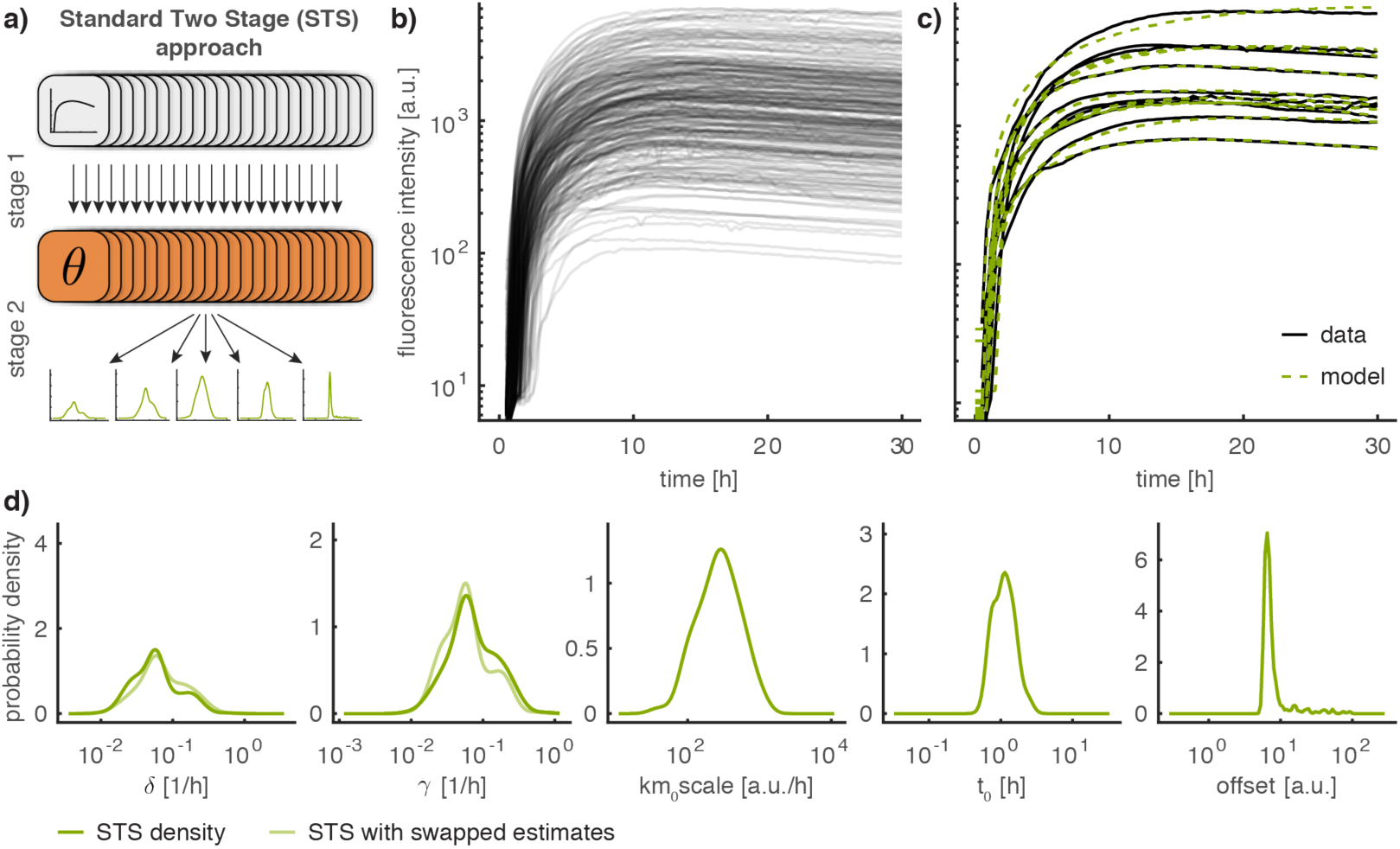
Parameter estimation results for the STS approach. **a)** Schematic illustration of the Standard Two-Stage (STS) approach. **b)** Experimentally recorded single-cell eGFP trajectories. **c)** Exemplary fits for 10 single-cell trajectories. **d)** Parameter distributions computed according to the STS approach using a kernel density estimate. The symmetry in *δ* and *γ* is illustrated by showing the respective kernel density estimate if the estimated values are swapped in lighter color.

The parameter distributions of the degradation rates, *δ* and *γ*, span two orders of magnitude and indicate multiple modes (Fig. 2d). At first glance, this suggests a previously unidentified subpopulation structure. Yet, the careful examination of the model structure (see Supplementary Information, Section *Uncertainty Analysis of Single-Cell Parameters*) revealed symmetry in the two degradation rates, *δ* and *γ*. We found that this symmetry, instead of multiple subpopulations, gives rise to the particular shapes of the estimated parameter distributions. This symmetry in the parameter estimates corresponds to a global structural non-identifiability (Chis et al. 2011).

Using structural identifiability analysis, we found that the symmetry could be resolved by (i) simultaneously measuring single-cell mRNA and protein levels or by (ii) measuring single-cell protein levels in cells sequentially transfected with different mRNA constructs. Both approaches are conceptually feasible (see (Chis et al. 2011; Darmanis et al. 2016; Frei et al. 2016) for (i)) but non-trivial. In general, the unique identification of relevant parameters from a single experiment is often challenging or impossible with the available experimental techniques. For bulk experiments, structural non-identifiabilities can sometimes be resolved by considering additional perturbation experiments (Raue, Kreutz, et al. 2013). This is possible as parameters can be assumed to be conserved quantities, which enables the recasting of perturbation experiments as additional observables in the model (Ligon et al. 2017). In contrast, it is non-trivial to consider additional perturbation experiments in the STS approach. For the STS approach, all single-cell parameters, and, hence, the parameter distributions for all experiments are assumed to be independent. Thus, no conserved quantities exist and information from multiple perturbation experiments cannot be exploited efficiently.

### Multi-Experiment NLME-Modeling Breaks Parameter Symmetry

To resolve the structural non-identifiability, we employed the NLME approach to integrate multiple perturbation experiments. In contrast to the STS approach, the NLME allows for conserved quantities between perturbation experiments (Fig. 3a). For the NLME model, we assume that the parameters *φ*_*i*_ of the *i*th single cell consist of a fixed effect *β* and a random effect *b*_*i*_ ∼ *N*(0, *D*): *φ*_*i*_ = exp(*β* + *b*_*i*_). The population parameters *β* and *D* are the mean and the covariance of the logarithm of the single-cell parameters. This corresponds to a lognormal distribution assumption, which ensures positivity of single-cell parameters.

**Figure 3.**
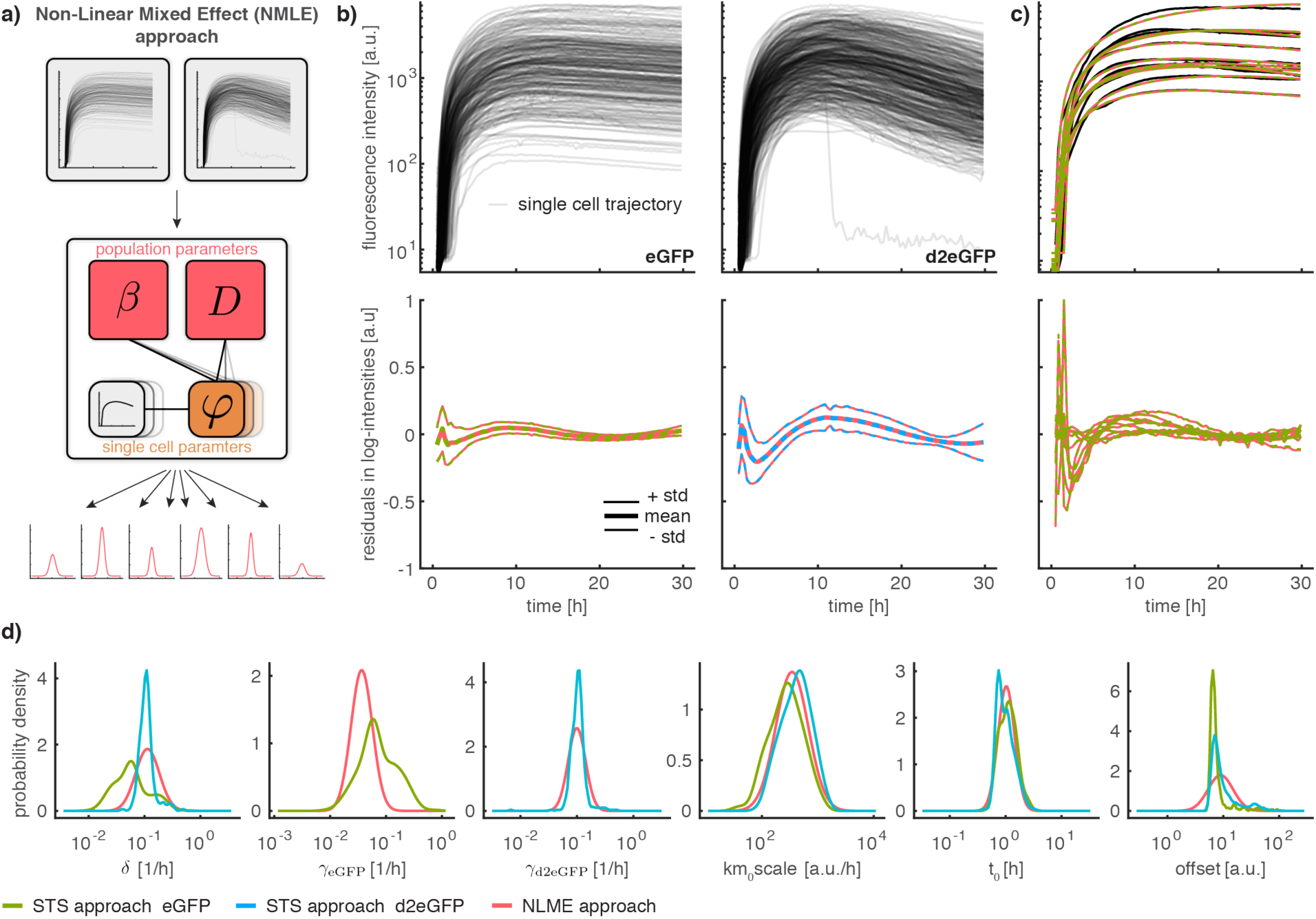
Comparison of parameter estimation results for the NLME and STS approaches. Coloring indicates employed approach (STS, NLME) and dataset (eGFP, d2EGFP). **a)** Schematic illustration of the NLME approach. **b)** Experimentally recorded single-cell eGFP and d2eGFP trajectories (top) and population statistics of residuals for the investigated approaches (bottom). **c)** Exemplary fits for 10 representative single-cell trajectories (top) and corresponding residuals (bottom). **d)** Comparison of parameter distributions computed using the STS and the NLME approach.

The population parameters *β* and *D* are conserved between experimental conditions, even if different individual cells are observed. Accordingly, single-cell data collected for different experiments can be integrated on the level of the cell population using *β* and *D*. In the NLME approach the population parameters *β* and *D* as well as the single-cell parameters *φ*_*i*_ are estimated simultaneously in a hierarchical optimization problem (see Supplementary Information, Section *NLME Approach*).

To resolve the non-identifiability in the translation model, we recorded an additional dataset in which cells were transfected with destabilized eGFP (d2eGFP). The fluorescence intensity was quantified for single cells for a duration of 30 hours after transfection. For cells transfected with eGFP the recorded signal reached a peak at around 9h and remained stable for the rest of the experiment (Fig. 3b). For cells transfected with d2eGFP the signal peaked around 10h and declined subsequently. For both datasets a pronounced variability of the recorded trajectories is evident.

We used both datasets, eGFP and d2deGFP, to estimate parameters using the STS and the NLME approach. For the NLME we assume two distinct parameter distributions for the protein degradation rates (*γ* _*eGFP*_ for eGFP and *γ* _*d2eGFP*_ for d2eGFP) (see Supplementary Information, Section *Multi-Experiment Extension of the NLME Approach*). For all other parameters, including the mRNA degradation rate *δ*, the distributions are assumed to be identical across experiments. For the STS approach, we also assume that the protein degradation rates differ and calculate the corresponding parameter distributions for each experiment. However, it is not possible to enforce that the distributions of the remaining parameters are identical between the experiments.

The analysis of the optimization results revealed that the STS and the NLME approach yield almost identical fits to the single-cell data (Fig. 3c). Moreover, for the NLME approach, the distributions of the estimated single-cell parameters *φ*_*i*_ agreed with the estimated distribution parameters *β* and *D* (Fig. S2). In summary, this suggests that the distribution assumptions for the NLME were appropriate.

For most of the parameters (*km*_0_*scale, t*_0_, *offset* and *γ* _*d2eGFP*_), both approaches yield similar population distributions (Fig. 3d). This may be surprising for *γ* _*d2eGFP*_, as the same symmetry effects as for *γ* _*eGFP*_ could be expected. However, the estimated degradation rates for mRNA and protein are so close to each other that symmetry has a negligible effect. For the mRNA degradation rate *δ*, the STS approach yields different distributions depending on the considered dataset, while the NLME, by construction, yields a single consistent distribution. For *γ* _*eGFP*_, the NLME yields a narrower distribution than the STS approach. This narrowing of the distribution can be attributed to the breaking of the parameter symmetry through the consideration of an additional dataset. (see Supplementary Information, Section *Uncertainty Analysis of Single-Cell Parameters*). This demonstrates that only the NLME approach is able to convey meaning to single-cell perturbation experiments. Indeed, by breaking the symmetry, the NLME approach improves not only the estimates of population parameters such as mean and variance, but also estimates of single-cell parameters (Fig. S1).

### Model Selection indicates Rate Limitation through Ribosome Abundance

To assess the appropriateness of the fitting results achieved using the STS and NLME approaches for modeling, we computed the distribution of the residuals at different time points. The result revealed a clear temporal trend (Fig. 3b), indicating that the considered mechanistic model was insufficient to describe the data. For this reason, we analysed three additional models (Fig. 4a (ii)-(iv)) that take the effect of ribosomes in protein translation and enzymes on mRNA degradation into account. These extensions are supported by experimental evidence collected in other studies (Parker & Song 2004). Yet, the relevance of these processes for single-cell transfection dynamics has not been studied in detail.

**Fig. 4.**
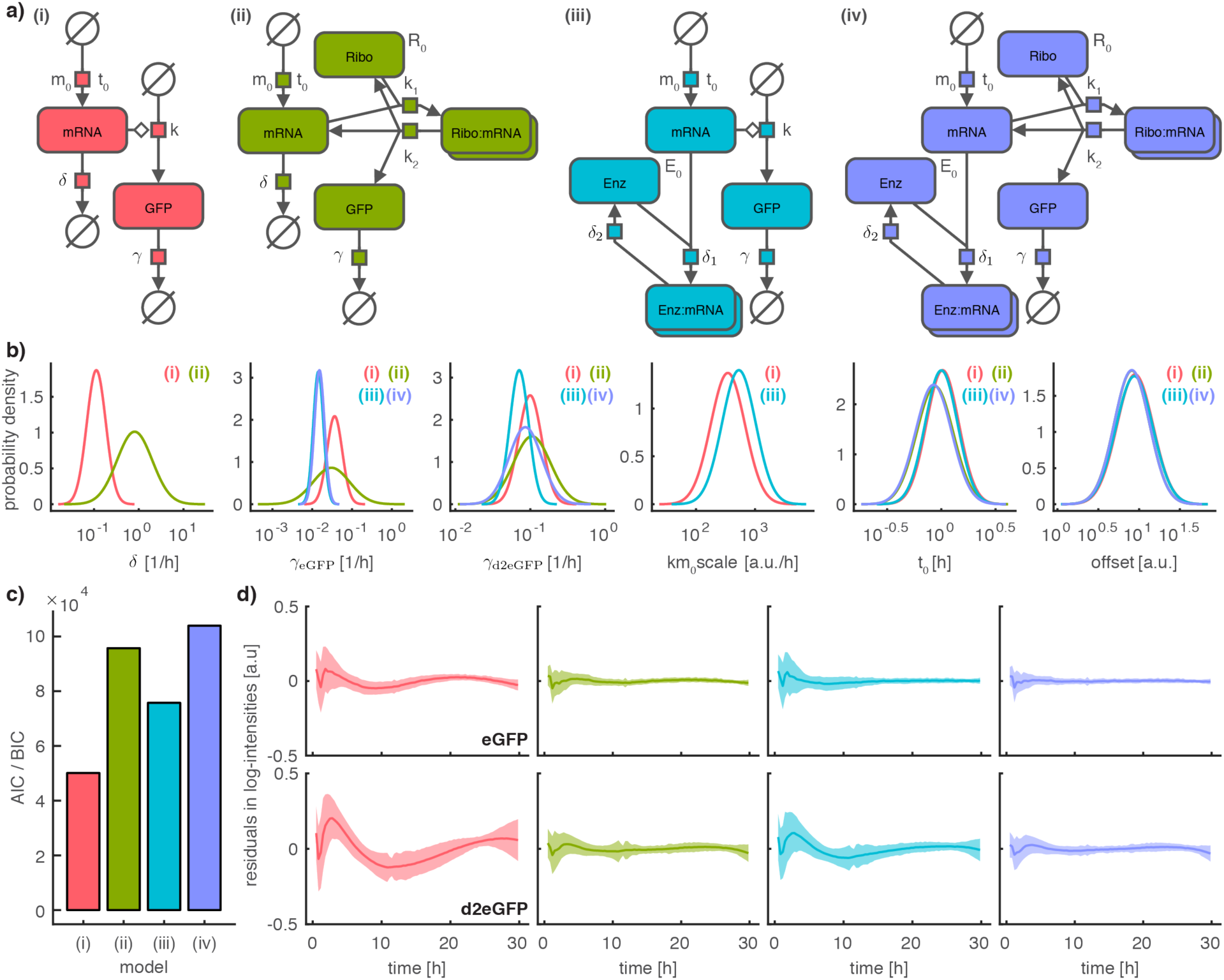
Comparison of parameter estimation results for four model candidates. Coloring corresponds to the model. **a)** Systems Biology Graphical Notation (SBGN) representation of the four model candidates with parameters plotted next to the respective reactions. The SBGN representation was created using the Newt editor (Sari et al. 2015). **b)** Comparison of parameter distributions across all considered models according to the NLME approach. Distributions are only shown for those models that include the respective parameter, as indicated by models numbers in each subplot. **c)** Comparison of log-likelihood values for MLE estimate for the four different models. **d)** Comparison of residuals for the four model candidates. Top: eGFP, bottom: d2eGFP. Shaded areas correspond to +/-one standard deviation across single-cells.

The effect of ribosomes is modeled by introducing the ribosomal species with initial free concentration *R*_0_, which binds to the mRNA at rate *k*_1_ (Fig. 4a (ii)). The translation happens at rate *k*_2_ and produces a free mRNA molecule and a free ribosome. The effect of enzymatic degradation is modeled by introducing an enzymatic species with initial free concentration. The degradation reaction of the mRNA is replaced by a binding reaction to the enzyme with rate *δ*_1_ (Fig. 4a (iii)). The enzyme degrades the bound mRNA molecule with rate *δ*_2_. In the standard model, these reactions are approximated by first order kinetics, which is reasonable under the assumption that the abundance of ribosomes and enzymes are not rate limiting. In total we considered four different models (Fig. 4a): **(i)** the standard model with two extensions that feature **(ii)** ribosomal binding to mRNA before translation as well as **(iii)** enzymatic degradation of mRNA and **(iv)** the combination of both extensions.

We inferred the parameters of models (i) - (iv) using the multi-experiment NLME approach. The corresponding optimization problem was solved using multi-start local, gradient-based search, which provided reproducible estimates (Fig. S3). The comparison of estimated parameters revealed striking differences (Fig. 4b), in particular for the degradation rates *δ, γ* _*eGFP*_ and *γ* _*d2eGFP*_. This highlights the need for appropriate models when estimating these kinetic parameters.

To select among candidate models, we considered the model selection criteria Akaike Information Criterion (AIC) (Akaike 1998) and Bayesian Information Criterion (BIC) (Schwarz 1978). AIC and BIC favored the two models with ribosomal translation (Fig. 4c), model (ii) and (iv). The best AIC and BIC values were achieved by model (iv) — the most complex model —, accounting for ribosomal translation and enzymatic degradation. Closer inspection revealed that the AIC and BIC values are dominated by the value of the log-likelihood function. The contribution of the complexity penalization is minor.

To confirm these findings, we evaluated the time-dependent residual distribution (Fig. 4d). This revealed that the magnitude of the residuals is substantially smaller for models with ribosomal translation (ii,iv) compared to models which do not account for this mechanism (i,iii). This suggests that the approximation of the ribosomal translation process with first order kinetics is not appropriate and that the abundance of free ribosomes is rate-limiting. In contrast, the difference in the residual profiles for model (ii) and (iv) is small, suggesting that enzymatic degradation is not essential to describe the data. For this reason, we perform all subsequent analyses with model (ii).

### Mechanistic Model Identifies and Explains Batch Effects

The experimental setup used in this study allowed for the investigation of multiple perturbation conditions in a single experimental batch. This is not always possible and each experimental condition often has to be investigated in an individual batch, which makes it difficult to distinguish perturbation specific effects from batch effects. Moreover, studies considering single-cell data sometimes pool batches to increase the statistical power (Zechner et al. 2014), which makes it difficult to distinguish single-cell heterogeneity from batch effects. Hence, robustness to batch effects is required for the meaningful integration of multiple (pooled) perturbation experiments.

To assess the robustness and reproducibility of our results, we recorded two additional experimental replicates, i.e., two additional batches. All replicates provide single-cell trajectories for >200 cells per experimental condition over a duration of ∼30 hours. For each experimental replicate, the single-cell trajectories were split into 3 subsets of same size, resulting in a total of 9 datasets (Fig. 5a). Even for our highly quantitative experimental setup, we found differences between replicates, which are larger than the sampling error (Figure 5b).

**Fig. 5.**
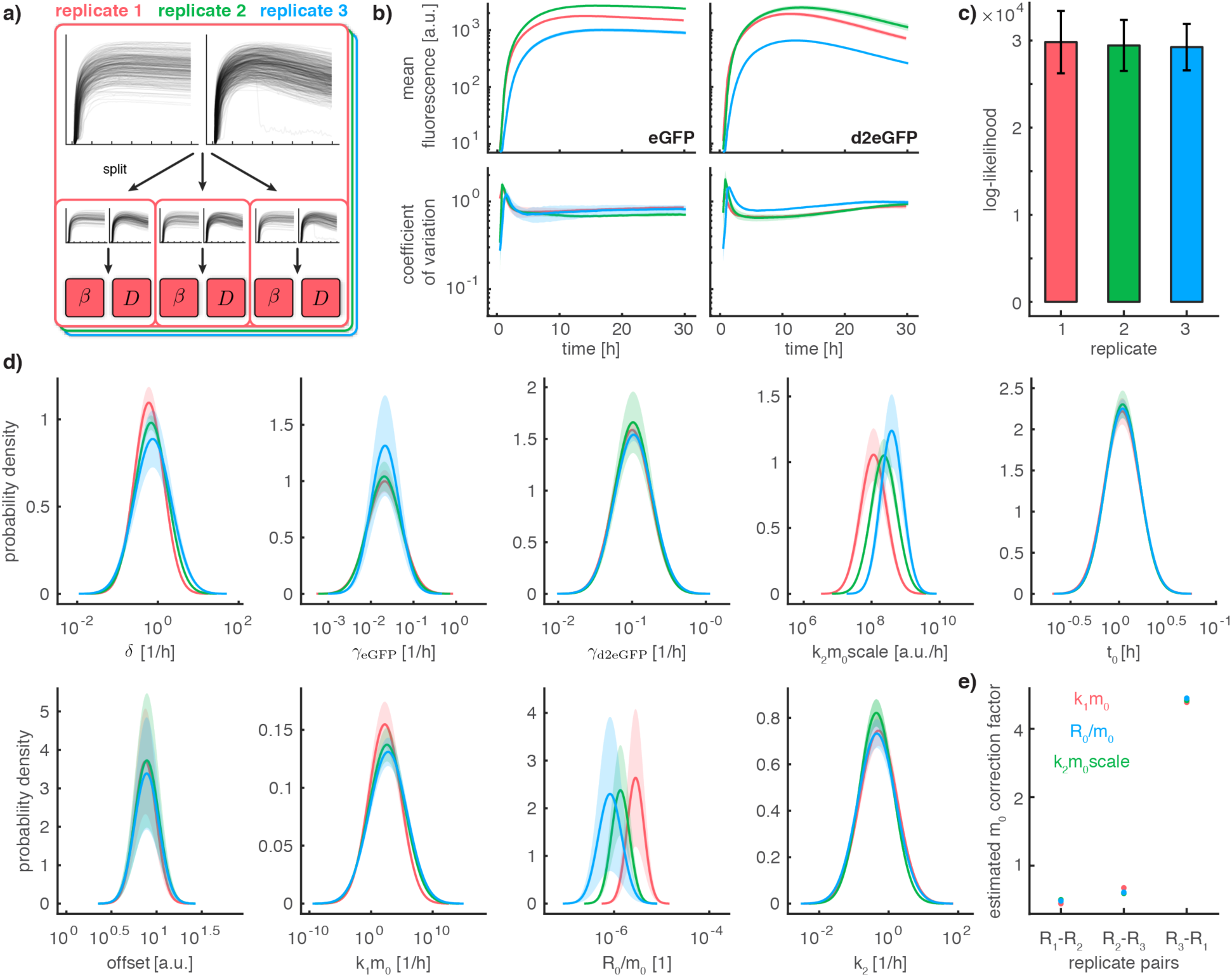
Uncertainty analysis for parameter distributions for model with ribosomal translation. Coloring according to replicate. **a)** Illustration of replicates and threefold split in data subsets **b)** Comparison of variability of mean and coefficient of variation within and across replicates. Shaded areas correspond to +/-one standard deviation within replicates. **c)** Comparison of log-likelihood within and across replicates. Error bars correspond to +/-one standard deviation within replicates. **d)** Analysis of uncertainty of parameter distributions within and across replicates. Shaded areas correspond to +/-one standard deviation within replicates. **e)** Estimated correction factors for m_0_ according to the respective estimated parameter distributions. Coloring according to the parameter distribution used to compute the correction factor.

To assess the effect of these batch effects on parameter estimates, we performed parameter estimation using the NLME approach for all 9 subsets individually. For all replicates, we observed similar average log-likelihood values, suggesting that the fit to all three datasets was of a similar quality (Fig. 5c). In contrast, we observed a much higher variability of log-likelihood values across the different subsets compared to individual replicates. As the log-likelihood is computed as a sum over all considered cells, the standard deviation can be rescaled to other settings by multiplying with the cell number ratio. This allowed us to compute the standard deviation of the log-likelihood values for the full experiment, which was considered for the analysis in Fig. 4c and accordingly test whether the observed difference in log-likelihood values for individual models is statistically significant. We found that the difference in log-likelihood value between model (i) and (ii) is statistically significant (p=0.018), while the differences between model (ii) and (iii) (p=0.084) and between model (ii) and (iv) (p=0.317) are not statistically significant.

The estimated parameter distributions were largely consistent within as well as across replicates (Fig. 5d). The only parameters for which we found apparent differences were *k*_2_*m*_0_*scale,k*_1_*m*_0_ and *R*_0_/*m*_0_ (due to the width of the densities for *k*_1_*m*_0_, the differences in *k*_1_*m*_0_ are hard to spot by eye). For these parameters, the population means differed substantially across replicates. All three parameter combinations contain the parameter *m*_0_, which describes the average concentration of mRNA transfected into the cells. However, since transfection efficiency is sensitive to cell culture conditions and the exact timing in the transfection preparation, it is likely that the experiment–to-experiment variability of the parameter *m*_0_ exceeds the intrinsic width of the parameter distributions of a single experiment. We found that the computed correction factors for *m*_0_ are consistent for all parameters and replicates (Fig. 5e). This suggests that differences in the average number of mRNA molecules released into the cytoplasm can indeed explain the observed batch effects, and shows that the NLME approach allows for the identification and handling of batch effects.

## Discussion

Single-cell time-lapse experiments are essential for the quantification and unraveling of the dynamics of cellular processes at the single-cell level. The NLME approach provides a powerful statistical tool to analyze single-cell fluorescence trajectories (Zechner et al. 2014; Karlsson et al. 2015; Almquist et al. 2015; Llamosi et al. 2016). In this study we introduced the multi-experiment NLME approach and demonstrated how it can be used to integrate single-cell measurements from multiple perturbation experiments. In particular, we studied eGFP expression after mRNA transfection using two distinct eGFP variants with different protein lifetime. The integration of these two datasets into a multi-experiment non-linear mixed effect modeling allowed for the robust parameter estimation. In particular, the reliable estimation of the distribution of mRNA lifetime, the key parameter of interest, is now possible for the standard model. Interestingly, the integration of data on the population level also improves the estimates of single-cell parameters by eliminating the structural non-identifiability (Fig. S1). The multi-experiment integration is an important advancement. Generally, experiments are measured using the same readouts under different experimental conditions, which essentially alters the entire set of parameters and does not change the quality of the parameter estimation.

The parameter estimation was performed using a gradient-based approach, which yields consistent results within as well as across replicate experiments, i.e., individual optimization runs consistently converged to similar likelihood values. Still, as observed in other studies (Kreutz 2016), we found that the convergence frequency of the local optimizers to the global optimum decreases as the model complexity increases (see Supplementary Information, Section *Reproducibility of Optimization Results for the NLME Approach*, Fig. S3). Moreover, although the current implementation of the approach scales linearly with respect to the number of considered single cells, application to higher throughput experiments may be problematic due to the already high computational cost for hundreds of cells.

The single-cell resolution allowed us to identify a pronounced temporal structure in the residuals of the established standard model of gene expression. This assessment revealed a substantial limitation of the established model and can directly be employed in other studies. Our model selection revealed improved agreement with a model that includes ribosomal rate-limitation of translation. Still, the residuals of the proposed model show a temporal structure and a higher magnitude for early time points (Fig. 4d). The experimental data show smoother onset dynamics than the model simulations (Fig. 4d, column 2), which could be explained by including GFP maturation or a more complex transfection process. The time-distributed delivery of multiple individual lipoplexes or a maturation phase of the protein would both be possible explanations for the smoother onset dynamics.

We find that for some of the parameters the population variability is estimated to be surprisingly large (e.g. *k*_1_*m*_0_ in Fig. 5). This may be due to a combination of large variabilities in *k*_1_ and *m*_0_. Alternatively, the description of the transfection or the translation process process might still be too simple, and that the variability accumating along multi-step processes is compensated by a large variability of few parameters (Snijder & Pelkmans 2011). As in similar studies, the results depend on the underlying mechanistic and statistical model. A refinement might improve the estimation accuracy further. Here, we see our results corroborated by the generally good agreement between literature values and the population mean parameter estimated with the NLME approach (Tab. S2).

All our findings were reproducible across independent experimental replicates. However, we found that batch effects arise even in sophisticated pipelines. In single-cell studies, these batch effects are often not handled adequately or hidden by pooling cells. Here, we demonstrated that the NLME approach can identify sources of batch effects, a crucial step to improve the experimental processes.

In summary, the NLME approach provides a powerful tool for the analysis of single-cell data with many possible extensions (Almquist et al. 2015; Karlsson et al. 2015; Kuhn & Lavielle 2005). We have demonstrated that the usefulness of the NLME approach is not limited to settings with sparse and noisy data, but also extends to settings where multiple experiments are considered. We anticipate that this unique feature of the NLME approach will help unraveling more mechanistic details about basic processes such as gene expression, but also enable careful design of targeted treatment strategies with mRNA-based therapeutic agents. Furthermore, we advocate the use of multiple datasets and their integration using the NLME approach to unravel underlying mechanisms and potential sources of batch effects.

## Author Contributions

The study was designed by A.R, F.F., F.T., J.H. and J.R.. A.R. performed all the experiments including image analysis. L.F. and D.W. developed the plug-in for background correction of the microscopy images. F.F. developed and performed the data analysis with help of L.F., J.H. and T.L.. A.R., F.F., J.H. and J.R. wrote the manuscript. The final version of the paper was commented and edited by all authors.

### Acknowledgments

We thank Mehrije Ferizi and Christian Plank for providing the two mRNA constructs used for this study. A.R. and F. F. are supported by a DFG Fellowship through the Graduate School of Quantitative Biosciences Munich (QBM).

## SUPPLEMENTARY INFORMATION

### Experimental Methods

#### Materials

RPMI 1640, Leibovitz’s L15 Medium, and Trypsin-EDTA were purchased from c.c.pro GmbH, Germany. FBS, HEPES solution, Sodium pyruvate, OptiMEM, and Lipofectamine™ 2000 were purchased from Invitrogen, Germany. Sterile PBS was prepared in-house. Six-channel slides (sticky μ-Slide VI^0.4^) and uncoated coverslips were purchased from ibidi, Germany. PLL(20kDa)-g[3.5]-PEG(2kDa) was purchased from SuSoS AG, Switzerland. Fibronectin was purchased from Yo Proteins, Sweden. For fluid handling on the microscope the selfmade tubing system was built of PTFE Microtubing with an inner diameter of 0.3 mm (Fisher Scientific, Germany), needlefree swabable valves (MEDNET, Germany), female luer lugs (MEDNET, Germany), and in-house made male luer teflon plugs.

#### Plasmid vectors and mRNA production

Open reading frame of Enhanced Green Fluorescent Protein (eGFP) as well as from Destabilized Enhanced Green Fluorescent Protein (d2eGFP) was excised from peGFP-N1 and pd2EGFP-N1 respectively (Clontech) and cloned into the backbone pVAX1-A120 which has been described previously (Kormann et al. 2011) to generate pVAXA120-eGFP as well as pVAXA120-d2EGFP.

The resulting plasmids were further linearized downstream of the poly(A) tail by NotI digestion and purified by chloroform extraction and ethanol precipitation. Purified linear plasmids were used as template for mRNA production via in vitro transcription (IVT) using RiboMax Large Scale RNA production System-T7 (Promega, Germany). Along with the addition of Anti-Reverse Cap Analog (ARCA) into to the IVT reaction mix to generate 5′ capped mRNA, chemically modified nucleotides, namely methyl-CTP and thio-UTP (Jena Bioscience, Germany), were added into reaction for the production of chemically modified mRNAs (cmRNAs) as described by Ferizi et al (Ferizi et al. 2015). The final RNA pellet was resuspended in RNAse free water and stored at −80°C.

#### Cell Culture

The human hepatoma epithelial cell line HuH7 (I.A.Z. Munich, Germany) was cultured in RPMI 1640 medium, supplemented with 10% FCS, 5mM HEPES and 5mM Sodium pyruvate. The cell line was cultured in a humidified atmosphere at 37°C and 5% CO_2_ level.

#### Surface patterning and array preparation

The single cell arrays are produced using microscale plasma-initiated protein patterning (μPIPP) on a polymer substrate as described in previous work (Ferizi et al. 2015; Segerer et al. 2016). In this study we used uncoated coverslips as substrate for the micropatterning which were glued to adhesive six-channel slides. Each bottom of the six channels is microstructured with the same pattern made of 30 μm × 30 μm adhesion squares, which are coated with Fibronectin, and a distance of 60 μm between the squares. The interspace between the Fibronectin squares is passivated with PLL(20kDa)-g[3.5]-PEG(2kDa).

HuH7 cells were seeded at a density of 10,000 cells per channel. The slide was stored in the incubator for four hours to enable cellular self-organization and adhesion on the Fibronectin squares (Röttgermann et al. 2014). Afterwards, two tubing systems each linking three of the six channels are connected with the slide enabling perfusion during the time-lapse measurement. The cell culture medium in the channels is exchanged to L15 medium containing 10% FBS by a syringe which is plugged in each of the swabable valves. The tubing systems were reused for all experiments after rinsing them with 70 % EtOH followed by distilled water and autoclaving them.

#### Time lapse microscopy

The time-lapse imaging was done on a motorized inverted microscope (Nikon, Eclipse Ti-E) equipped with an objective lens (CFI PlanFluor DL-10x, Phase 1, N.A. 0.30; Nikon). A heating chamber (ibidi GmbH, Germany) was used for controlling the sample temperature to 37 °C (± 2 °C) during the measurement. For image acquisition we used a cooled CCD camera (CLARA-E, Andor), a LED light source (SOLA SE II, lumencor), and a filter cube with the filter set 41 024 (CHROMA Technology Corp., Bp450-490, FT510, LP 510-565).

The six-channel slide connected with the tubing systems is put in the heating chamber and all tubes are fixed on the microscope stage for the time lapse measurement. The scanning macro containing information such as the exposure time as well as the position list is defined prior to the time-lapse measurement using NIS-Elements Advanced Research software (Nikon). The image acquisition was started right after adding the lipoplex solution to the cells. Fluorescent images were recorded every 10 min over a duration of 30 h.

#### In vitro Transfection on the microscope

The cells were transfected with Lipofectamine™ 2000 complexes containing one of the described mRNA constructs. Both mRNA constructs were treated with Lipofectamine™ 2000 using the following protocol: The lipoplexes are made of a ratio of 2.5 μl Lipofectamine™ 2000 per 1μg mRNA. mRNA and Lipofectamine™ 2000 were diluted separately in OptiMEM in a final volume of 250 μl each and incubated for 5 min at room temperature. The mRNA solution and the Lipofectamine™ 2000 were mixed and further incubated for 20 min at room temperature for lipoplex formation. The final lipoplex solutions have a total volume of 500 μl with a mRNA concentration of 0.5 ng/μl.

During the lipoplex formation the cells were washed with 37 °C warmed-up PBS. After the incubation each of the tubing systems were rinsed with 500 μl of one of the mRNA lipoplex solutions. The time point when the lipoplex solution was added to the cells is defined as the beginning of the experiment. After 1 h of lipoplex incubation the cells were washed with 37 °C warmed-up L15 medium supplemented with 10% FBS and the cells were not treated further for the remaining measurement.

### Mathematical Methods

#### Data Acquisition and Quantitative Image Analysis

The obtained raw image stacks of the time-lapse experiments were processed using in-house written plugins in ImageJ for background correction and readout of the single-cell trajectories (Ferizi et al. 2015). First, the image stacks were background corrected based on a previously published algorithm (Schwarzfischer et al. 2011). Afterwards, the fluorescence trajectories were generated by calculating the mean intensity over time for every cell on the micropattern, which expressed the fluorescent protein. To read out the fluorescence trajectories, a grid corresponding to the micropattern is aligned to every image stack and only squares occupied by a cell are manually selected to calculate the mean intensity of the square for every frame of the image stack.

Analysis of processed data suggested a constant offset and multiplicative measurement noise in the recorded fluorescence trajectories. Therefore, we log-transformed the data, which yields additive noise for which standard algorithms can be used. The log-transformation is also applied to observable map in the models, which will be described in the following section. The model also includes a scaling and offset parameter, which is estimated along with the other model parameters.

#### Mathematical Models for GFP Translation

In the following we provide model equations for all four considered models of GFP transfection. For all models we initially derived a basic form of the model and then identified structurally non-identifiable parameters using the MATLAB toolbox GenSSI (Ligon et al. 2017). Subsequently, we applied state transformation and conservation laws to eliminate structural non-identifiabilities and unnecessary state equations. The model simulations were all carried out using the AMICI toolbox (Fröhlich, Kaltenbacher, et al. 2017; Fröhlich, Theis, et al. 2017). We provide AMICI and SBML implementations of the model in Supplementary File S2. The ODEs for model (i) - (iv) are provided in Table S1.

The models consider up to four state variables: mRNA abundance (*m*), protein abundance (*p*), ribosome abundance (*r*), enzyme abundance (*e*), ribosome-mRNA complex abundance (*rm*) and enzyme-mRNA complex abundance (*em*). All models include the bolus injection which increases the mRNA level at time *t*_0_ from 0 to *m*_0_,

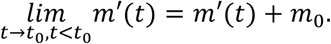

In the transformed models, we consider the normalized mRNA abundance, *m* = *m*′/*m*_0_, yielding for the bolus injection

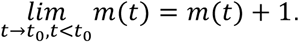

The bolus injection is not explicitly described in Table S1. Instead, we write *m*′(*t*_0_) = *m*_0_ and *m*′(*t*_0_) = 1. The mRNA abundances are 0 up to time *t*_0_.

**Table S1.**
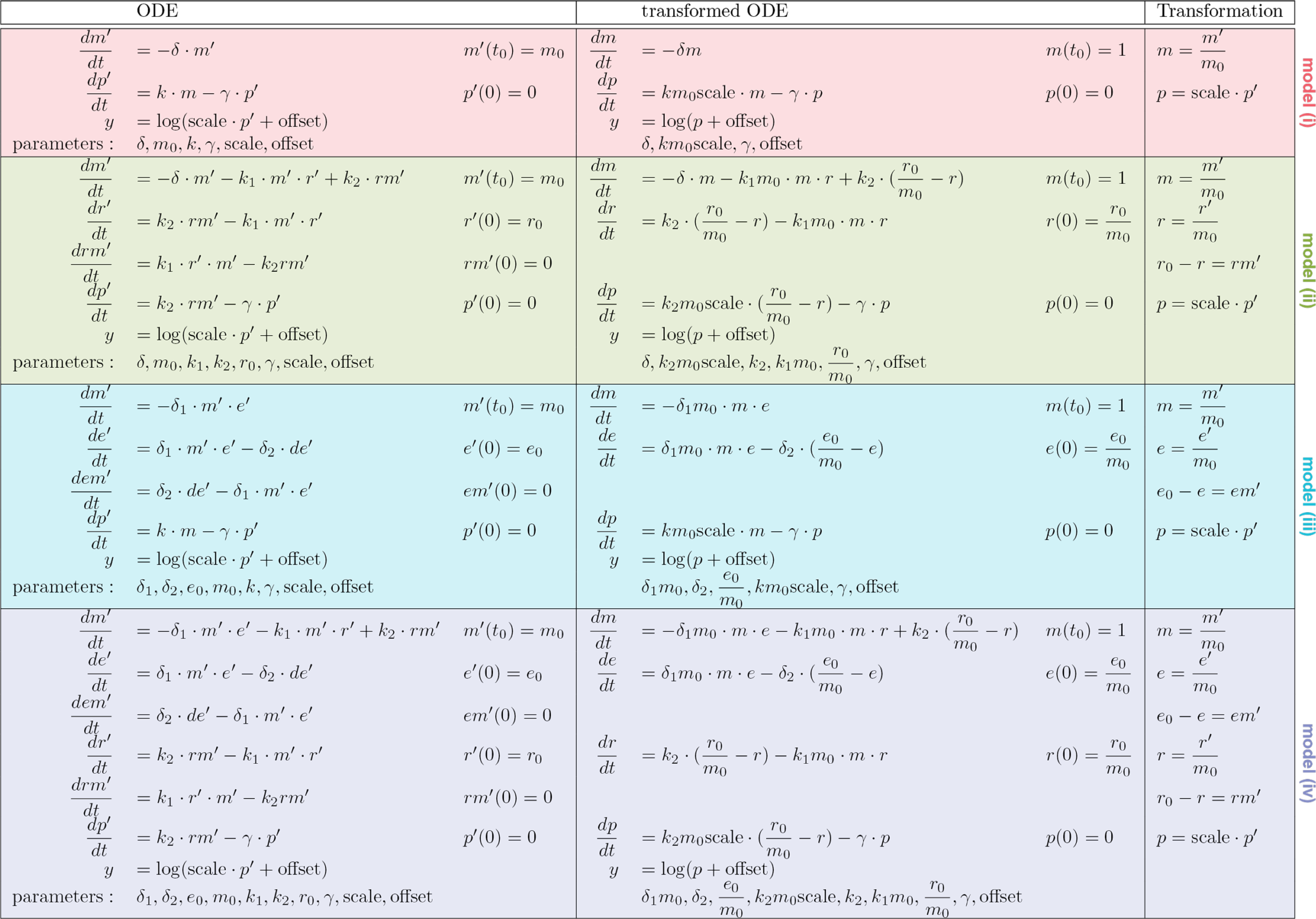
Mathematical Formulation and Transformation of Models (i) - (iv). Molecule abundances are abbreviated for mRNA (m), protein (p), ribosome (r), enzyme (e), ribosome-mRNA complex (rm) and enzyme-mRNA complex (em).

#### Uncertainty Analysis of Single-Cell Parameters

To assess the effect of the population constraint on single-cell parameter estimates we analyzed the respective uncertainty for the STS and the NLME approach using the profile likelihood approach (Raue, Schilling, et al. 2013). The analysis revealed that the uncertainty of parameters *km*_0_*scale, t*_0_ and *offset* is negligible compared to the population variability (Fig. S1a). Moreover, there are only small differences in the location of the mode and the width of the profiles between the two approaches, which suggests that the imposition of the population constraint in the NLME approach has negligible effect on the estimated values for these parameters. For *δ* and *γ* _*eGFP*_ the profiles for the STS approach are bimodal, which can be expected due to the parameter symmetry. For the single-experiment (eGFP dataset) NLME approach, the separation of modes increased compared to the STS approach, but parameter symmetry persisted. For the multi-experiment NLME approach, the profiles become unimodal, which indicates that the symmetry has indeed been broken.

A careful inspection of profiles revealed that a large number of profiles of *δ* and *γ* _*eGFP*_ are unimodal instead of bimodal. This effect can be explained by the superposition of uncertainty of the two modes that would be expected according to parameter symmetry. To demonstrate this effect we approximated the profile shown in Fig S1a as sum of two normal densities (Fig. S1b). For the investigated case, a change in the mean of the normal densities, as small as 1% of the parameter value, led to the emergence of a single mode in the middle between the two previous modes. As *δ* and *γ* _*eGFP*_ are symmetric, a unimodal profile implies that both parameters are estimated to the same value. Consequently, the STS approach fails to resolve sufficiently small differences between *δ* and *γ* _*eGFP*_ and instead estimates both parameters to be the same (Fig. S1c). In this study, this leads to a trimodal shape of the parameter population density (Fig. 1d), where two of the modes are explained by the symmetry of parameters for which the difference was sufficiently big, while the the third mode is explained by the remaining parameter for which the difference could not be resolved. Overall, this phenomenon introduces a bias in the estimates of the single-cell parameters as well as the population level distributions.

For the NLME approach, we do not directly observe this effect, as all profiles are unimodal. However, we found that some of the single-cell profiles are slightly skewed. This might be the reason for the slight skewness in the estimated density of single-cell parameters (Fig S1d). Yet, as the NLME approach assumes a normal density, this skewness is not present in the estimated population variability.

**Figure S1.**
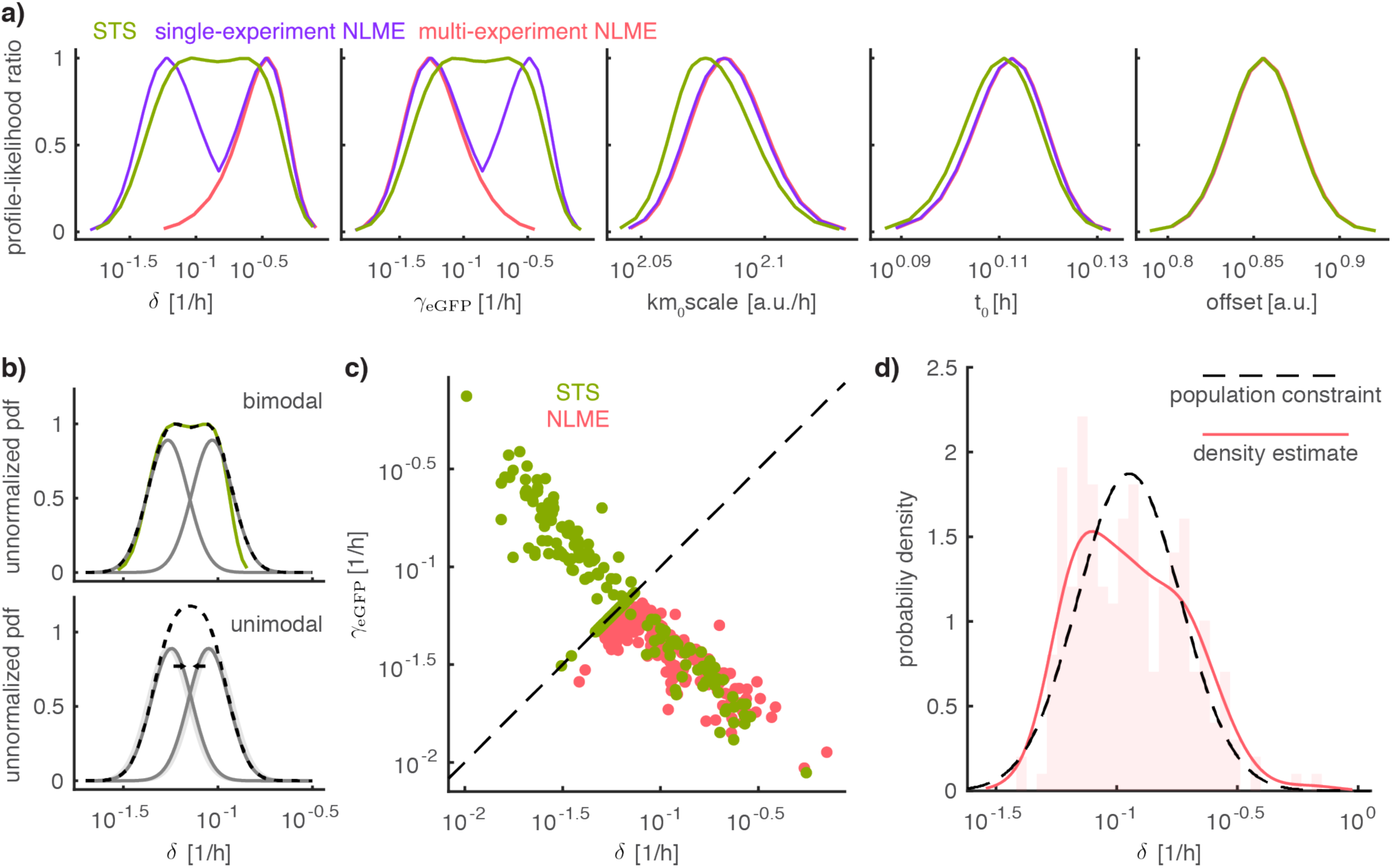
Analysis of single-cell parameters for the STS and the NLME approach for the eGFP dataset for a single cell. **a)** Profile-Likelihood ratios of single-cell parameters for the STS approach (*θ*_*i*_, green), the single-experiment NLME approach (*φ*_*i*_, purple) and for the multi-experiment NLME approach (*φ*_*i*_, red). For the NLME approach profiles, the estimated population parameters *β* and *D* were fixed during profile calculation. For the single-experiment NLME approach profiles from local minima in the parameters *β* and *D* were combined. Color indicates the employed approach. **b)** Sketch explaining the emergence of unimodal and bimodal profile shapes. The dashed black line is the superposition of the two normal densities in grey. The computed profile-likelihood is shown as green line. **c)** Comparison of estimated single-cell parameters for the STS (*θ*_*i*_, green) and NLME (*θ*_*i*_, green) approach. **d)** Comparison of NLME single-cell parameters histogram (bars) and kernel density estimate (red line) with the population parameter distribution (black dashed line).

**Figure S2.**
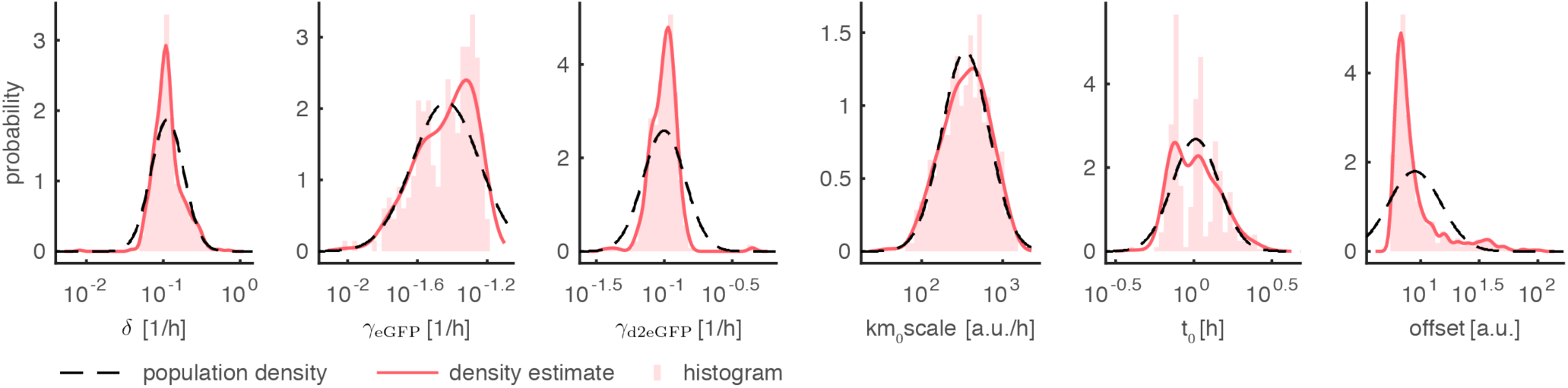
Comparison of distributions of estimated single-cell parameters (*φ*_*i*_) and estimated population parameter distributions (*β*,*D*). NLME single-cell parameters histogram (bars) and kernel density estimate (red line) with the population constraint (black dashed line).

**Figure S3.**
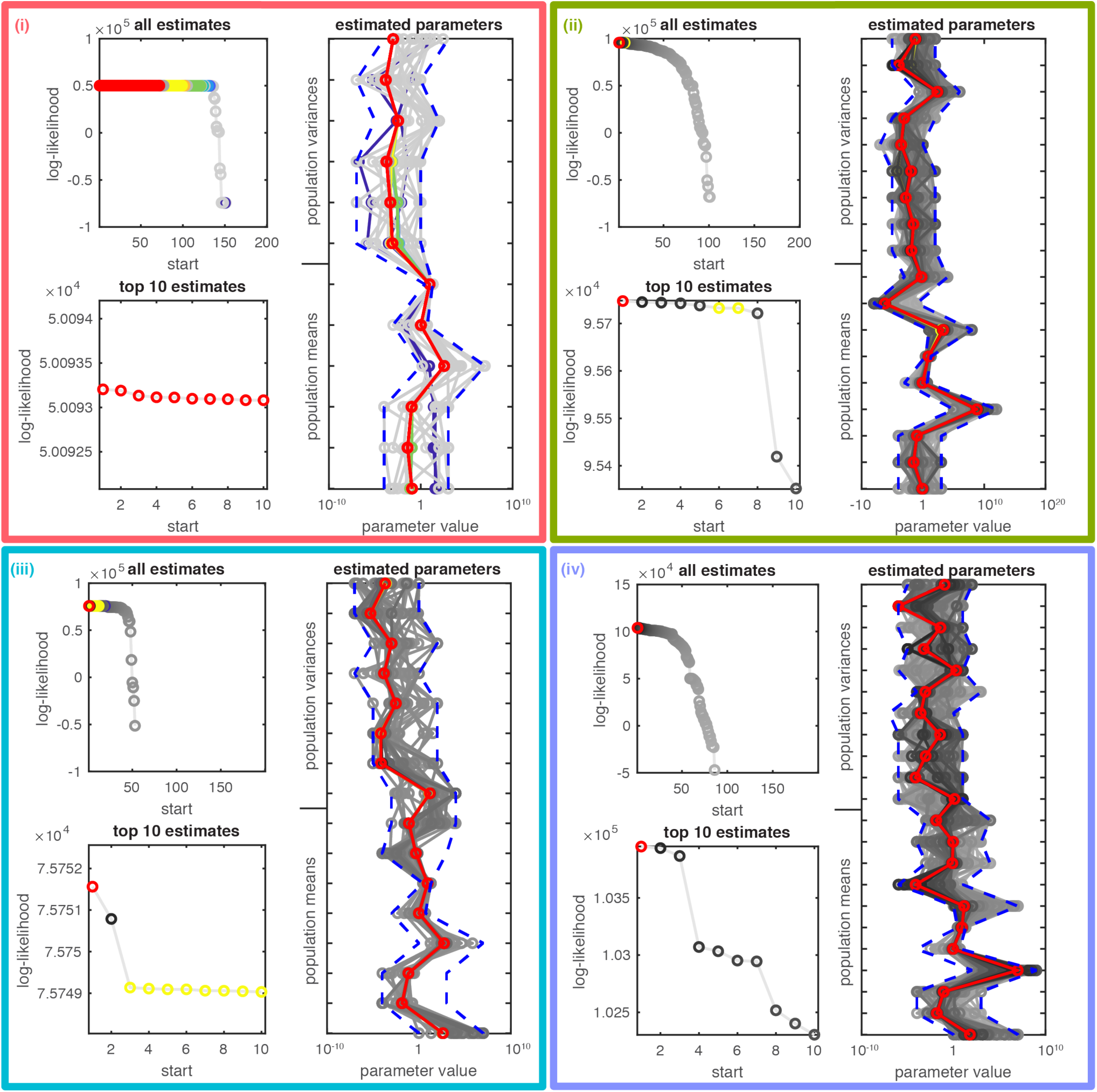
Analysis of reproducibility of optimization results for the NLME approach. Results for each considered model are shown in individual subplots, which are marked by roman numerals and border color. In each subplot, the left two plots show all/top10 log-likelihood values across all optimization runs. Highest objective function value is always colored in red. For all other optimization runs, coloring is only applied if multiple optimization runs yielded objective function values within 0.01 of each other, otherwise a grey color indicates the rank in log-likelihood value. The same coloring is applied to the estimated parameter values on the right. Crashed optimization runs are not shown.

#### Literature Validation of Estimated Parameter Values

Our structural identifiability analysis revealed that only few of the kinetic rates could be uniquely estimated as for most only the products of rates were identifiable. For the uniquely identifiable mRNA degradation rate *δ* and the protein degradation rates of *γ* _*eGFP*_ and *γ* _*ed2GFP*_ we were able to compare the estimated values to literature values from bulk experiments (Table S2).

**Table S2.** Comparison of estimated population mean values to literature values for the NLME approach with model (ii).

| Parameter | Formula | Estimated value | Literature value | Reference |
| --- | --- | --- | --- | --- |
| mean mRNA (all genes) half-life | $\frac{\ln(2)}{\delta}$ | 0.8h | 1-30h | (Schwanhäusser et al. 2011) |
| mean eGFP half-life | $\frac{\ln(2)}{\gamma_{eGFP}}$ | 22.8h | ~26h | (Corish & Tyler-Smith 1999) |
| mean d2eGFP half-life | $\frac{\ln(2)}{\gamma_{d2eGFP}}$ | 6.6h | ~5.5h | (Corish & Tyler-Smith 1999) |

#### STS Approach

For the STS approach we used a standard Maximum Likelihood method to estimate parameters. We transformed the data and assumed additive, independent, normally distributed measurement noise with variance 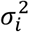. This yields the likelihood function

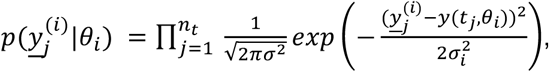

where 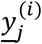 is the measurement data for cell *i* and the vector *θ*_*i*_ contains the unknown single-cell model parameters. In a single-cell context, no empirical estimator, which requires multiple replicates, can be used to estimate 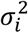 from data, as we are limited to a single replicate per single cell. Therefore, we use the model-based estimate of 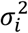 (Pinheiro 1994):

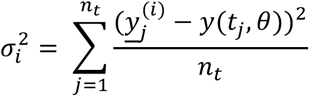

To estimate the parameters *θ*_*i*_, the optimization problem

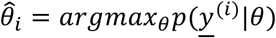

is solved for every cell using a local gradient-based multi-start algorithm implemented in the Matlab toolbox PESTO (Stapor et al. 2017). AMICI (Fröhlich, Kaltenbacher, et al. 2017) was used for the computation of the gradient. The convergence was checked by comparing the best estimates. The population densities were then computed based on a kernel density estimate.

#### NLME Approach

For the NLME approach we replace the parameter *θ*_*i*_ by a mixed effect *φ*_*i*_ which consists of a fixed effect *β* and a random effect *b*_*i*_ ∼ *N*(0, *D*). To compute the likelihood for the NLME model, we have to evaluate an integral over the random effects (Pinheiro 1994):

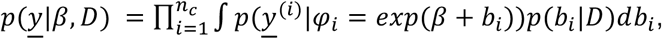

where *n*_*c*_ is the total number of cells. Numerically exact evaluation of this integral with methods such as Markov Chain Monte Carlo sampling is computationally highly demanding. Instead, we used the Laplace approximation (Pinheiro 1994). The Laplace approximation assumes that 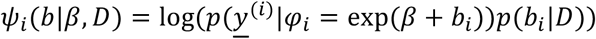 is close to a quadratic function in *b*, which allows analytic evaluation of the integral for the approximate likelihood *p*:

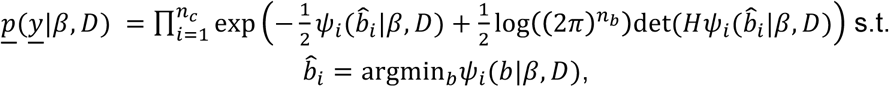

where 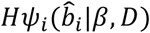 is the Hessian of 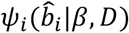. To estimate the population parameters with the NLME approach, we solved a hierarchical optimization problem with the outer problem

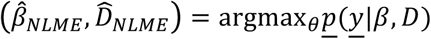

and the inner problem

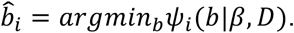

The computation of *<u>p</u>*(*y*|*β*, *D*) including the respective inner optimization problem was solved using the MATLAB toolbox MEMOIR (https://github.com/ICB-DCM/MEMOIR). In contrast to other NLME modeling toolboxes such as MONOLIX or NONMEM, MEMOIR employs local-gradient based optimization algorithm which uses AMICI (Fröhlich, Kaltenbacher, et al. 2017) for the computation of the gradient. The outer optimization problem was solved using Matlab toolbox PESTO (Stapor et al. 2017), using the interior-point algorithm implemented in FMINCON, initialized and 200 random starting points. MEMOIR was evaluated using a range of artificial data, which are included as examples in the repository.

For the covariance matrix, we used a Matrix Logarithm parameterization (Pinheiro 1994). The Matrix Logarithm parameterization guarantees that *D* is symmetric and positive definite and only uses a minimal number of parameters to prevent overparameterization. For the purpose of this study, we assumed that *D* has diagonal form.

#### Multi-Experiment Extension of the NLME Approach

The NLME approach *per se*, is not directly applicable to multiple experiments. We extended the approach by introducing experiment specific single-cell parameters 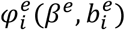 The corresponding likelihood function has to be computed separately per experiment

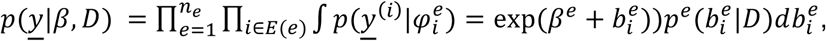

where *n*_*e*_ is the number of experiments, *E*(*e*) denotes the set of indices of cells belonging to a particular experiment and *p*^*e*^ is the density of the experiment specific population constraint corresponding to 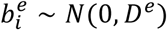. Without loss of generality, it is reasonable to assume that a single cell cannot belong to more than one experiment. The same approximation as before can be applied.

To encode the assumption of conserved quantities, we assume that parameterization of some of the parameters in *β* ^*e*^ and *D*^*e*^ are shared across the experiment. For this particular study, we assumed that the parameterization for all parameters, but *γ* _*eGFP*_ and *γ* _*d2eGFP*_, is shared across both experiments. For example for model (i), this means that the entries in *γ* ^*eGFP*^ correspond to the parameters [*δ, γ* _*eGFP*_, *km*_0_*scale, t*_0_, *offset*] and in *β* ^*d2eGFP*^ correspond to the parameters [*δ, γ* _*eGFP*_, *km*_0_*scale, t*_0_, *offset*].

#### Reproducibility of Optimization Results for the NLME Approach

For the NLME approach, we started 200 optimization runs per model. Each individual optimization run required 1-8 weeks of computation time (>15 years of cpu time in total). Our analysis revealed that the optimization results for model (i) were highly reproducible, as the log-likelihood values for the top 10 optimization runs had low variance. Moreover, only a small fraction of optimization runs crashed, i.e. had repeated failures in the evaluation of the log-likelihood function due to numerical integration problems. For models (ii) - (iv) the variance in log-likelihood value as well as the number of crashed optimization runs is substantially higher compared to model (i). However, we found that the differences among the top 10 optimization runs for each model were substantially smaller (10^1^ to 5 ⋅ 10^2^) than differences in log-likelihood values across models (>*E*; 5 ⋅ 10^3^). Hence, we do not expect any negative impact on the validity of model selection.

